# Fluctuation-driven plasticity allows for flexible rewiring of neuronal assemblies

**DOI:** 10.1101/2022.03.01.482464

**Authors:** Federico Devalle, Alex Roxin

## Abstract

Synaptic connections in neuronal circuits are modulated by pre- and post-synaptic spiking activity. Heuristic models of this process of synaptic plasticity can provide excellent fits to results from *in-vitro* experiments in which pre- and post-synaptic spiking is varied in a controlled fashion. However, the plasticity rules inferred from fitting such data are inevitably unstable, in that given constant pre- and post-synaptic activity the synapse will either fully potentiate or depress. This instability can be held in check by adding additional mechanisms, such as homeostasis. Here we consider an alternative scenario in which the plasticity rule itself is stable. When this is the case, net potentiation or depression only occur when pre- and post-synaptic activity vary in time, e.g. when driven by time-varying inputs. We study how the features of such inputs shape the recurrent synaptic connections in models of neuronal circuits. In the case of oscillatory inputs, the resulting structure is strongly affected by the phase relationship between drive to different neurons. In large networks, distributed phases tend to lead to hierarchical clustering. Our results may be of relevance for understanding the effect of sensory-driven inputs, which are by nature time-varying, on synaptic plasticity, and hence on learning and memory.

## I. INTRODUCTION

Learning occurs through changes in the synaptic weights between cells in neuronal circuits. Understanding learning therefore requires working out the rules by which these changes happen at a single synapse, and then studying the consequence of this process at the network level. One major finding regarding the rules underlying synaptic plasticity was the observation that the synaptic weight change could depend on the relative timing of pre- and post-synaptic spikes [1, 9]. Theoretical work has studied how such spike-timing dependent plasticity (STDP) can shape recurrent connections in neuronal networks [2, 5] and serve to encode fixed point attractors in large-scale spiking models [18]. However, like all Hebbian plasticity rules, STDP is intrinsically unstable, and leads either to full saturation of all synaptic connections, or complete depression, depending on the sign of the integral of the plasticity window. Essentially, plasticity always occurs if the product of pre- and post-synaptic rates is non-zero, even if constant. Therefore, additional stabilizing mechanisms are required in order to allow for the emergence of non-trivial connectivity patterns [7, 13, 17, 18]. A multiplicative STDP rule, for which potentiation is progressively weaker the stronger the synapse, stabilizes weights but does not readily allow for the emergence of non-trivial network structure [10, 12, 15].

Here we consider the scenario in which plasticity only occurs in the presence of timevarying rates. This mode of plasticity seems particularly relevant for learning given that most salient events unfold over time and would be expected to result in time-varying firing rates in the relevant brain circuits. For simplicity we explore this plasticity regime by assuming that the integral of the STDP window is exactly zero, i.e. that there is a balance between potentiation and depression when firing rates are constant. This obviates the need for added stabilizing mechanisms and allows for an in-depth analysis.

We consider both oscillatory as well as noisy drive and develop a theoretical framework for pairs of linear firing rate neurons. Specifically, we make use of the separation of timescales between neuronal and synaptic dynamics to derive self-consistent evolution equations for the synaptic weights. Analysis of these equations reveals a rich phase diagram from which the resulting connectivity motif can be predicted depending on the phase difference in the case of oscillatory drive, or the correlation and delay in the case of noisy drive. We also find many regions of multistability, meaning that the final connectivity motif will also depend on the initial configuration of weights. For the case of oscillatory drive we study the effect of the balanced STDP rule in networks numerically, and show that the resulting connectivity matrix can be well predicted from the pairwise theory in several relevant cases.

## II. GENERAL TWO-NEURON MODEL

### A. Firing rate formulation for two neurons with forcing

We begin by considering the simplest possible scenario of two coupled neurons. We model the neuronal activity with a linear firing rate equation, which will allow for a complete analysis. The equations are

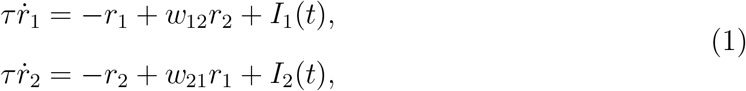

where *w*_12_ and *w*_21_ are the recurrent synaptic weights. The external inputs vary in time; we will consider both oscillatory inputs as well as correlated noise sources in subsequent sections.

### B. Plasticity Rule

Here we interpret the rates in Eqs.1 as the underlying probability for a Poisson spiking process, and then apply a so-called spike-timing dependent plasticity (STDP) rule [8]. That is, in a small time interval Δ*t*, the probability that neuron 1 generates a spike is just *r*_1_(*t*)Δ*t*. Once the spikes have been generated, a presynaptic spike of neuron *i* followed by a postsynaptic spike of neuron *j* at a latency *T* leads to a potentiation of synapse *w_ji_* by an amount *A*_+_*e*^−*T*/*τ*_+_^ and a depression of synapse *w_ij_* by an amount -*A_e*^−*T*/*τ*_−_^. Synapses are bounded below by zero and above by a maximum value *w_max_*. Whenever there is a new spike the synaptic weights are updated in this way for all past spike pairs, see [11] for an efficient numerical implementation. This rule, together with Eqs.1, provide a complete model which can be simulated numerically. Note that while we are formally making use of an STDP rule here, the exact spike timing plays no role. That is, plasticity is due only to dynamics in the rate. Such rates effects appear to dominate over contributions due to spike timing in models of STDP when realistic input patterns are considered. Specifically, experimental protocols have traditionally used highly regular, repeated pairings of pre- and post-synaptic activity, and the observed synaptic plasticity has therefore been natually attributed to the exact spike timing [1, 9]. However, theoretical work has shown that when the inferred rules are used in the presence of more *in-vivo* like spike trains with a high level of irregularity, the resulting synaptic plasticty can be accounted for to a large extent just by variations in the underlying firing rate [6].

If the amplitude of potentiations and depressions is small compared to the maximum synaptic strength *w_max_*, then the evolution of the synaptic weights can be approximated by the following integrals [8]

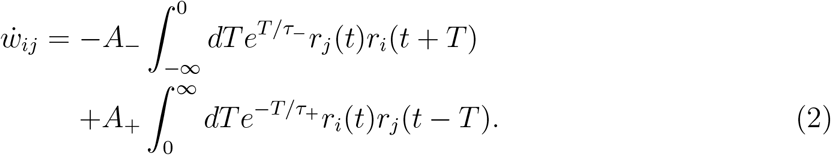

The first integral includes the contribution of all spike pairs leading to depression of the synapse *w_ij_*. Specifically, given Poisson processes, the probability of the spike pair in which cell *i* has spiked in a small interval around time *t* and cell *j* has spiked previously at a latency *T* is just *r_i_*(*t*)*r_j_*(*t* – *T*) (*T* > 0). The integral sums up spike pairs at all possible latencies up until the current time. The second integral is analogous, but for spike pairs which lead to potentiation of the synapse. When the neuronal dynamics is stationary, the integrals can be written in the compact form

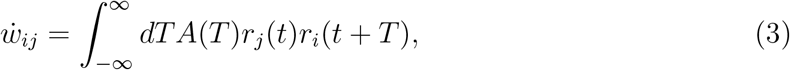

where *A*(*T*) = *A*_+_*e*^−*T*/*τ*_+_^ for *T* > 0 and −*A_e*^*T*/*τ*_−_^ for *T* < 0, see *Appendix A* for a detailed explanation. Eqs.3 together with the rate equations Eqs.1 consitute a self-consistent approximation to the full model, and which is ammenable to analysis.

### C. Asymptotic approximation for slow plasticity

Eqs.1 and 3 cannot be solved for directly. The reason is the presence of quadratic nonlinearities of the firing rate variables in the integrals in Eqs.3. However, we can take advantage of the slowness of synaptic plasticity compared to the firng rate dynamics to derive an approximate system of equations which can be solved exactly. Specifically, we assume that the amplitude of potentiation and that of depression are small and formalize this by replacing the kernel *A*(*T*) = *ϵĀ*(*T*) in Eqs.3. We also define a new, slow time *t_s_* = *ϵt* and allow the rates and the synaptic weights to evolve both on a fast as well as on a slow timescale. We expand the rates and weights in orders of e and find that the leading order solution, where 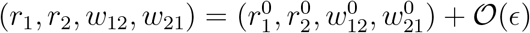, obeys the following coupled equations

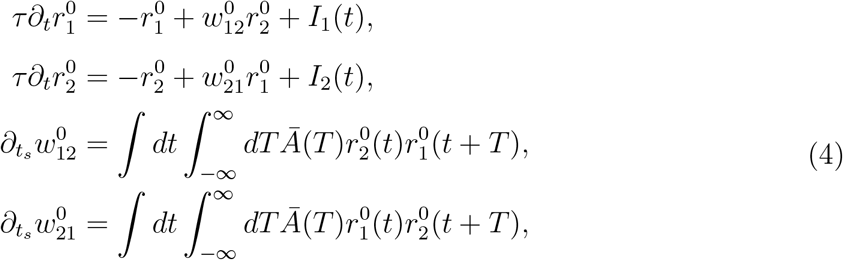

see *Appendix A* for details of the derivation. In Eqs.4 there is a formal separation of the timescale of evolution of the rates from that of the synaptic weights, which is much slower. In fact, the synaptic weights are only a function of the slow-time and hence can be treated as constants in the first two equations, allowing one to solve for the rates using techniques from linear algebra. The integrals can then be formally evaluated, yielding self-consistent evolution equations for the synaptic weights alone. The integrals over the fast time are performed over an appropriate time window, e.g. over one period of oscillation for oscillatory drive.

## III. OSCILLATORY DRIVE

We first consider the case of oscillatory drive with frequency *ω* and phase difference *ϕ*. Specifically, we take *I*_1_(*t*) = *Ie*^*iωt*+*iϕ*_1_^ and *I*_2_(*t*) = *Ie*^*iωt*+*iϕ*_2_^. The physiological firing rates are given by the real parts of *r*_1_ and *r*_2_, which are then also used to calculate the weights self-consistently in Eqs.4, yielding

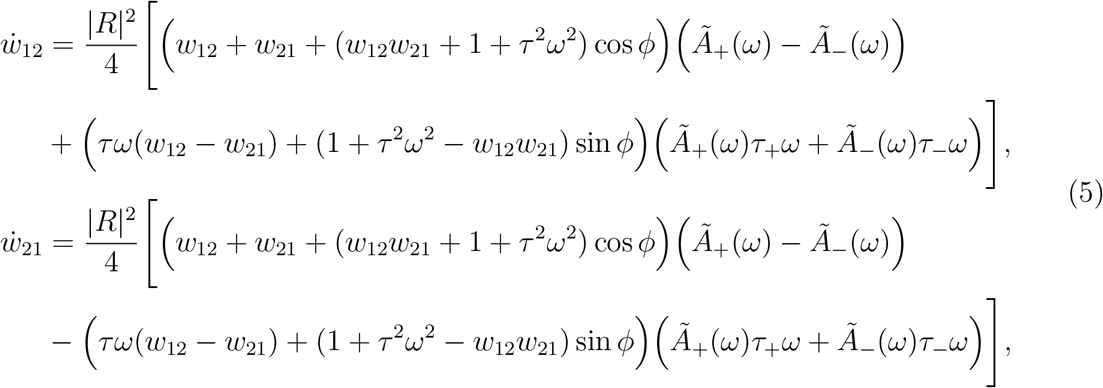

where 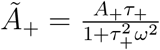, *ϕ* = *ϕ*_2_ – *ϕ*_1_ and we have left off the superscript 0 for simplicity.

An analysis of Eqs.5 reveals that there are no fixed point solutions for *w*_12_, *w*_21_ ≥ 0. However, by studying the sign of the right hand side in Eqs.5 at the boundaries of the allowable domains, we can find stable solutions. For example, the fully potentiated solution (bidirectional motif) is stable if 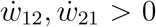 for (*w*_12_, *w*_21_) = (*w_max_, w_max_*). This condition is clearly satisfied for *ϕ* = 0 as long as *Ä*_+_(*ω*) – *Ä*_(*ω*) > 0, which holds when potentiation dominates at short latencies. On the other hand, when the *ϕ* = *π*, 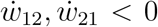, meaning that there is a critical value of the phase for which the fully potentiated solution becomes unstable. In a plane of the phase versus the frequency of the forcing, there is therefore a curve below which the potentiated solution is stable, see the orange line in Fig.1. An analogous argument can be made for the fully depressed solution (unconnected motif), which in this case is stable above a critical curve, see the blue line in Fig.1. Finally, the solution for which one synapse is fully potentiated and the other fully depressed (unidirectional motif) is stable between the dashed, black lines in Fig.1. Note that there is a region of bistability between the unidirectional motif and the bidirectional (unconnected) motif, indicated by the orange (blue) hatching in Fig.1.

**FIG. 1.**
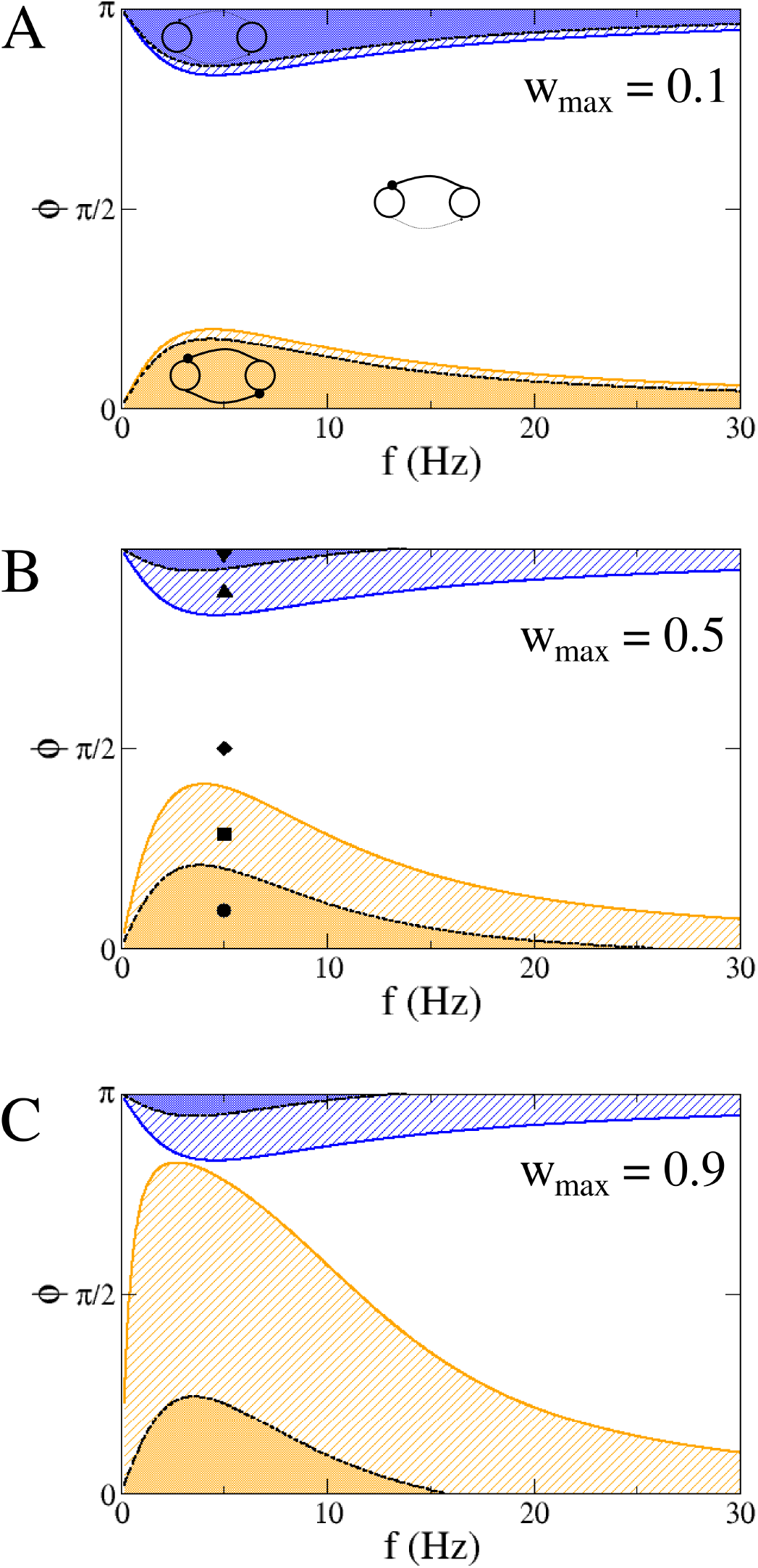
Phase planes of the synaptic dynamics in Eqs.5 as a function of the phase difference *ϕ* and frequency *f* = *ω*/(2*π*) of the forcing, and for different values of *w_max_*. A. Phase plane for *w_max_* = 0.1. Bidirectional, unidirectional and unconnected motifs are stable in the orange, white and blue regions respectively. The unidirectional motif is stable in the region demarked by the two dashed lines; hence the system is bistable in the hatched regions. B. Phase plane for *w_max_* = 0.5. The symbols indicate the parameter values used for the simulations in Fig.3. C. Phase plane for *w_max_* = 0.9. Other parameters: *τ* = 10ms, *τ*_+_ = 20ms, *τ*_−_ = 60ms, *A*_+_ = 0.001, *A*_−_ = *A*_+_*τ*_+_/*τ*_−_, *I*_0_ = 30Hz, *I* = 20Hz.

The phase planes shown in Fig.1 show a clear resonance for values of a critical forcing frequency, approximately around 5Hz in this case. Specifically, the fully potentiated and fully depressed states are stable over a much wider range of forcing phases in this regime. Additionally, and as illustrated in Fig.2, the rate of change of the synaptic weights is also maximal around this value of the frequency, and decreases to zero for zero frequency and in the limit of high frequencies. Both effects can be understood as the interaction of the forcing frequency with the window of plasticity, i.e. there is a “best” frequency which maximizes the integral in Eq.3. Precisely this resonance mechanism has been invoked to explain the role of theta oscillations in driving plasticity in rodent hippocampus [14]. This optimal frequency can be found by taking the derivative of the growth rate as a function of the forcing frequency. Doing so in the case where *w*_12_ = *w*_21_ = 0 leads to the simple relation 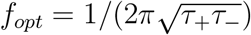, independent of forcing frquency, see vertical dashed line in Fig.2.

**FIG. 2.**
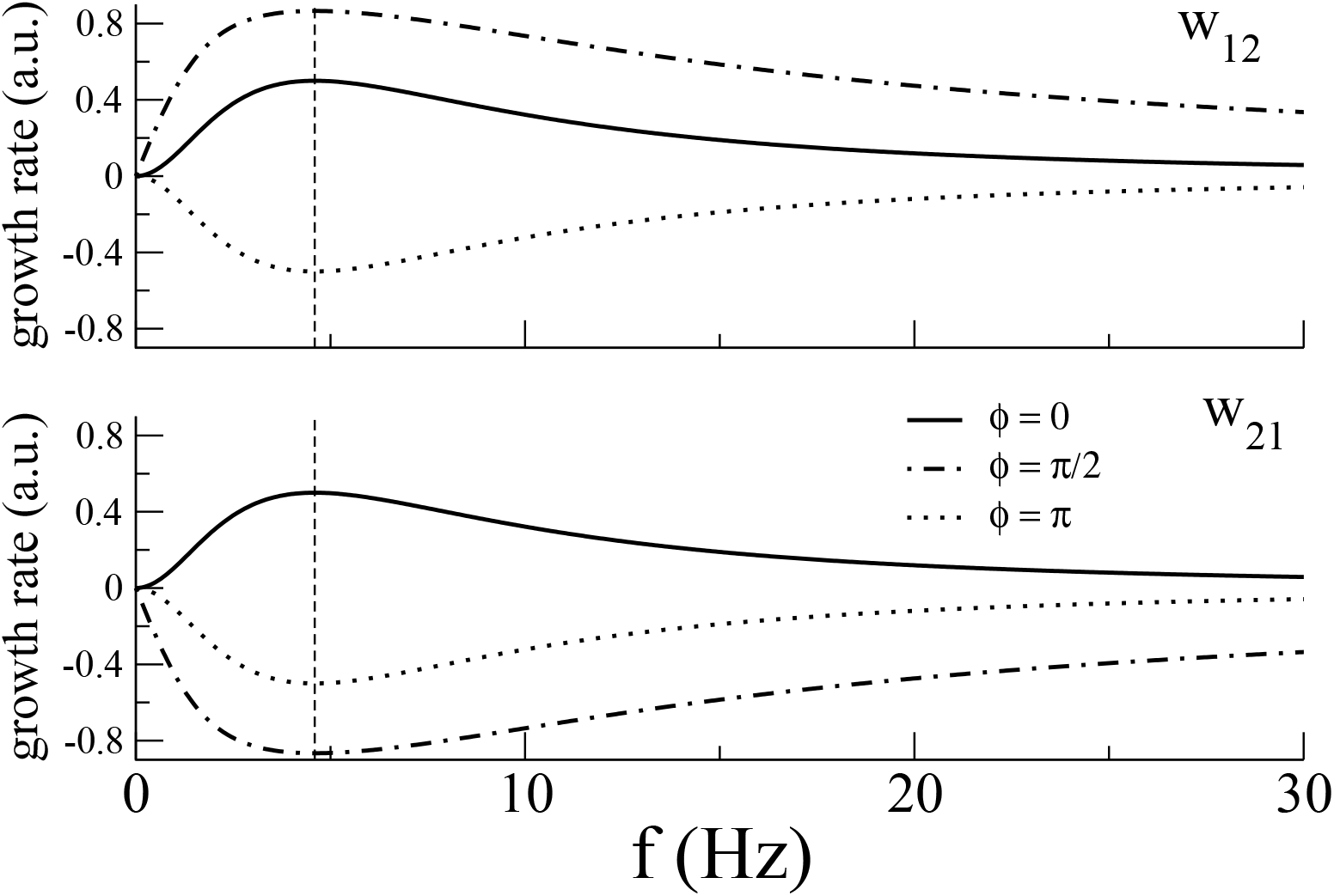
Growth rates proportional to 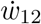 and 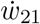 for (*w*_12_, *w*_21_) = (0,0) as a function of the forcing frequency and for three different phases *ϕ*. Note that the optimal forcing frequency is independent of phase and equal to 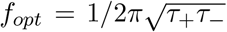 (vertical dashed line). All other parameters are as in Fig.1

Numerical simulations of the full model agree well with the analysis of the reduced system. Specifically, given a fixed forcing frequency, there is a range of phase-differences near the in-phase forcing which lead to both synapses potentiating (PP) (orange region in Fig.1). Fig.3 shows an example of the synaptic dynamics in this region for forcing frequency and phase indicated by the black circle in Fig.1B. For slightly larger phase differences two sets of synaptic weights can coexist depending on initial conditions: PP or one potentiated and the other depressed (DP) (orange hatched region in Fig.1), see example simulations in Fig.3 for the parameters given by the black square. In an intermediate range of phase differences between in-phase and anti-phase, only uni-directional connectivity emerges (DP) (white region in Fig.1), see an example simulation in Fig.3 for the parameters given by the black diamond. Finally, close to an anti-phase forcing there is a region of bistability between (DP) and a fully disconnected motif (DD), followed by a region in which only the DD solution is stable (blue hatched region and solid blue region in Fig.1 respectively), see sample simulations for the black up- and down-triangles in Fig.3 respectively.

**FIG. 3.**
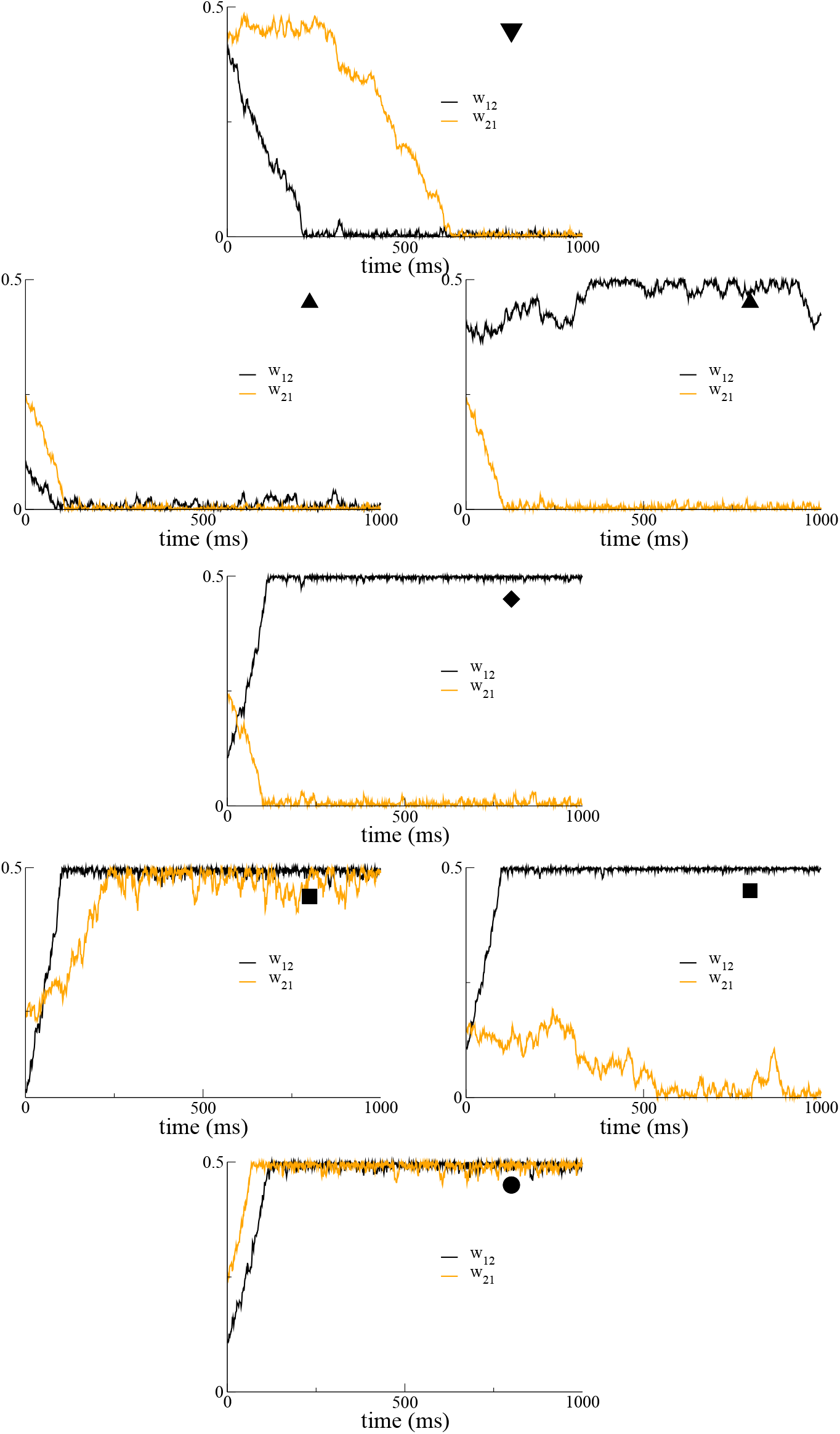
Sample synaptic dynamics for parameter values used in Fig.1B. The forcing frequency is fixed at *f* = 5Hz and the phase is indicated by the corresponding symbol in Fig.1. The different final states in the traces for identical parameter values are due to different initial conditions.

### A. Oscillation-driven plasticity in networks

In the previous section we derived evolution equations for the synaptic weights for a single pair of neurons. We showed that these equations can admit several different stable configurations of weights depending mainly on the phase difference of the forcing, while the frequency chiefly affected the learning rates. We now study the evolution of synaptic weights in a network of an arbitrary number of neurons. Once again we drive all of the neurons with an oscillatory forcing at a fixed frequency, and with a phase which can differ from cell to cell. The resulting synaptic weight matrix is expected to depend on the precise choice of phases. In principle we can make use of the same theory to derive a set of coupled ODEs for the synaptic weights, akin to Eqs.5, see *Appendix B* for details. This leads to N(N-1) coupled equations for a network of size N. However, there are some simple cases for which the resulting synaptic weight matrix can be straightforwardly predicted from the pairwise theory. The simplest example is that of a network in which half of the neurons are driven at one phase, and the other half at a different phase. In this case, neurons within a cluster have zero phase difference between them, leading to the strengthening of recurrent connections, while neurons from different clusters will have their connection shaped according to the given phase difference. For a phase difference of π, the between-cluster connections decay to zero, leading to the formation of two unconnected clusters, as shown in Fig.4B.

**FIG. 4.**
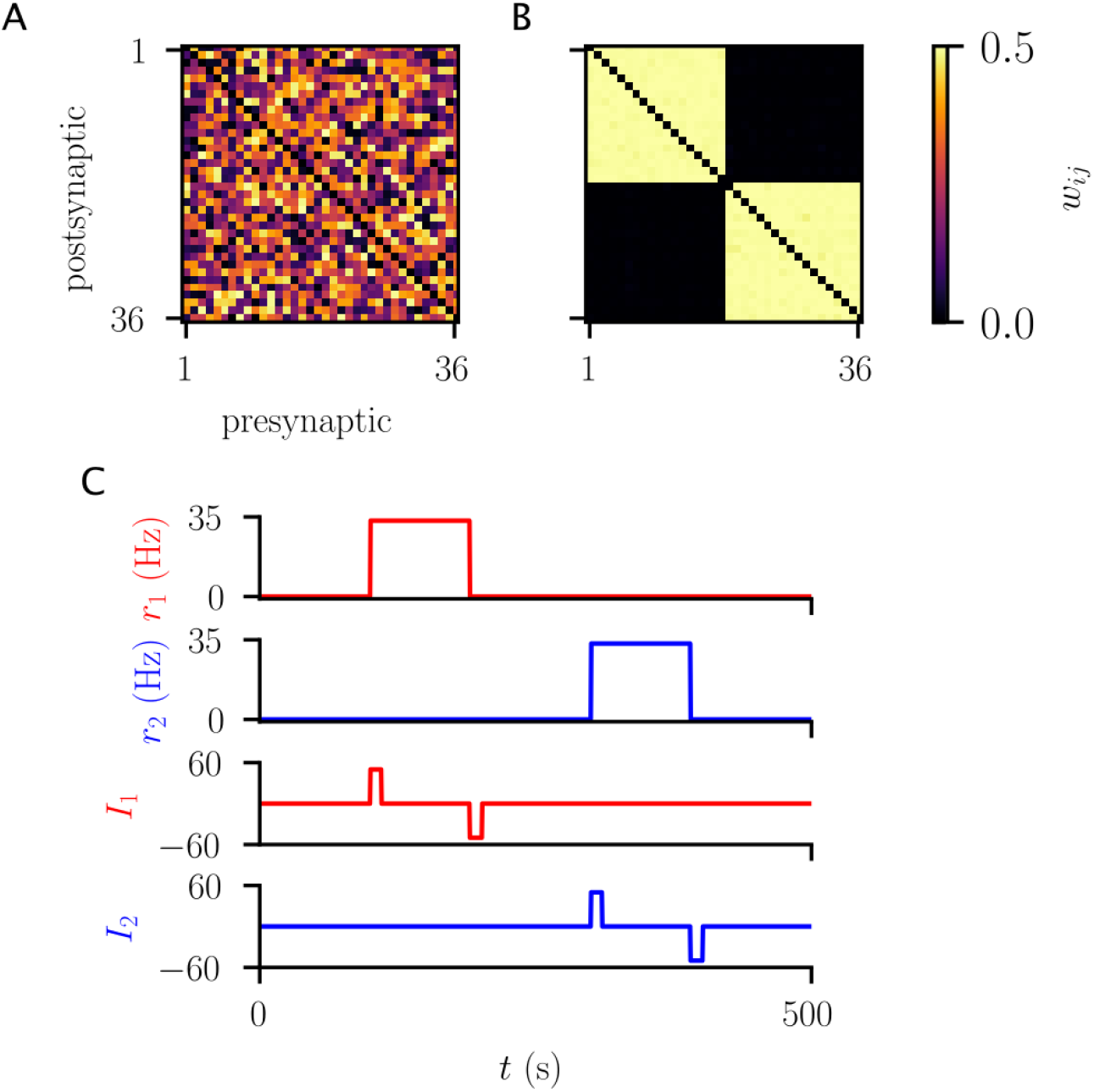
Driving two populations of neurons out-of-phase leads to the formation of two disconnected clusters. A. The initial synaptic weight distribution. Weights are uniformly distributed between 0 and 0.5. B. The synaptic weight matrix at long times after oscillatory forcing of two groups of 18 neurons each, with a phase difference of π between the two groups. Simulation time is 10^6^ms. Parameters: *I*_0_ = 5, *α*^−1^ = 0.03, *I* = *I_osc_* cos *ωt, I_osc_* = 30. Coupling normalization is *K* = *N_cl_*. C. After the learning period the two neuronal populations exhibit bistability due to the strong recurrent connections. For these simulations the neurons are taken to have a Heaviside transfer function.

Note that, even though the theory was developed for linear firing rate neurons, it nonetheless correctly predicts the final state of the weight matrix even for nonlinear rate neurons, at least in this simple case. In fact, for the simulations shown in Fig.4, the neuronal transfer function is taken to be a Heaviside function, see *Appendix B* for details. Given this choice, the clusters, once formed, can exhibit bistability, see Fig.4C.

The applicability of the pairwise theory to networks in which the forcing is clustered is further illustrated in Fig.5. Specifically, with a phase difference of *π*/2 in a two-cluster network, we expect the cross-cluster connectivity to be unidirectional, from the leader to the follower. This is indeed what is found in simulation, see Fig.5A. For the case of three clusters, in which clusters 2 and 3 have a phase difference of *π*/2 and *π* with cluster 1 respectively, Fig.5B shows that the same unidirectional motif is found between clusters 2 and 1, and 3 and 2, while 1 and 3 become uncoupled, all as predicted from the pairwise theory.

**FIG. 5.**
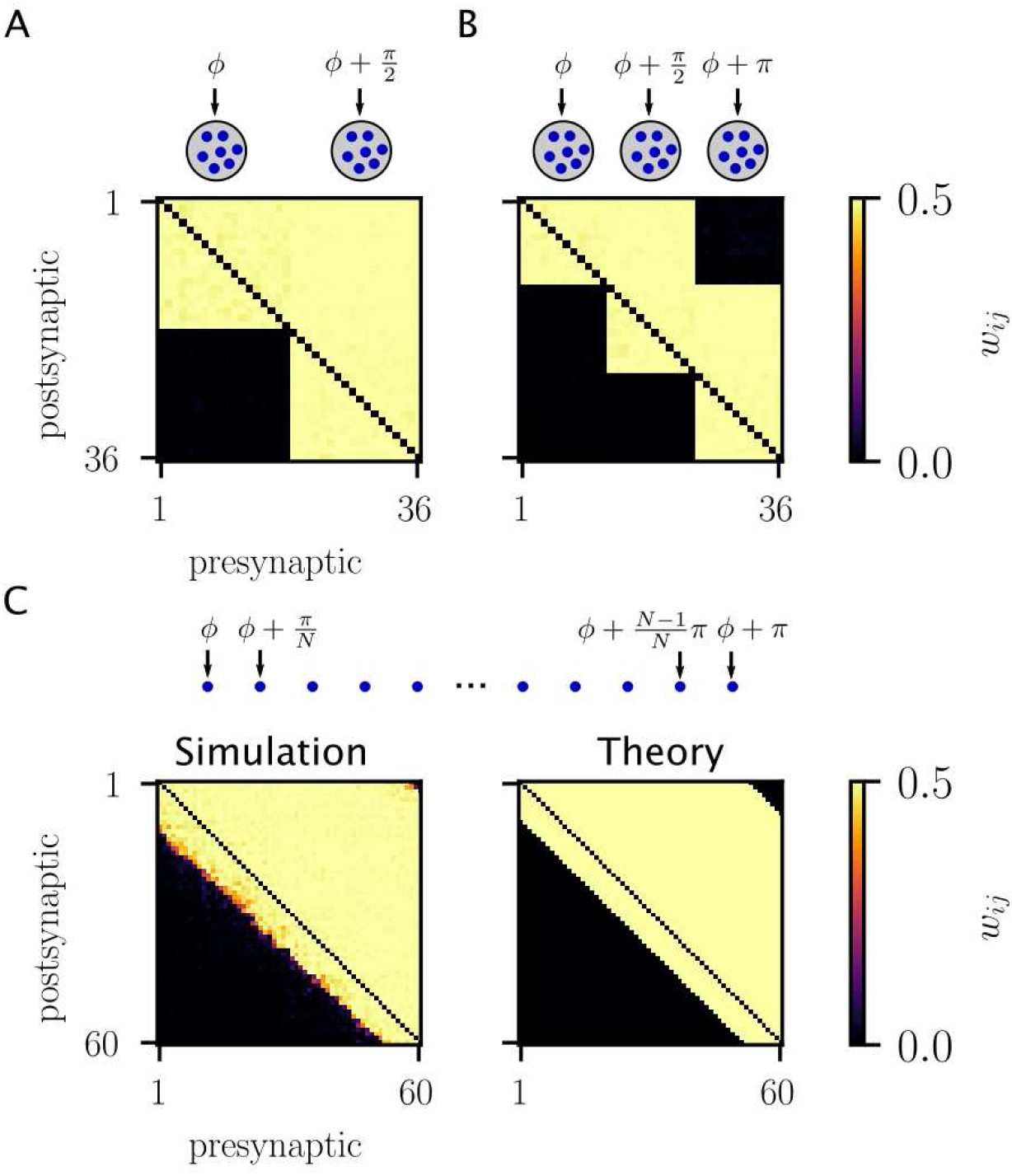
Network connectivity depends on the distribution of phases and is well-predicted from pairwise theory. A. A two-cluster network with a phase difference of *π*/2. B. A three-cluster network with phases *ϕ*_1_ = *ϕ, ϕ*_2_ = *ϕ* + *π*/2 and *ϕ*_3_ = *ϕ* + *π*. C. A network with phases distributed uniformly between 0 and *π*. Other parameters are the same as in Fig.4

The theory can furthermore be extended to the case in which the phase difference is distributed uniformly across the network, from 0 to *π*. In this case the interaction between any pair of neurons depends mainly on the weights between those two neurons because the influence of the rest of the network is close to zero, see *Appendix B* for details. In this case, neurons with similar phases are expected to form strong recurrent connections, while sufficiently different phase difference will lead to leader-follower unidirectional motifs, and phase differences near *π* will lead to complete uncoupling. Numerical simulations show that the resulting synaptic weight matrix is, in fact, very close to that predicted from the pure pairwise theory, see Fig.5C.

## IV. NOISY DRIVE

In this section we consider a pair of neurons driven by noisy inputs. The inputs are

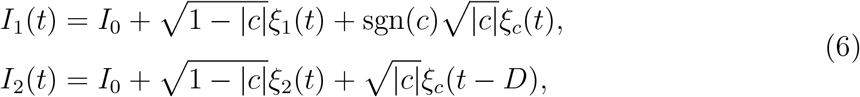

where 〈*ξ_i_*(*t*)〉 = 0 and 〈*ξ_i_*(*t*)*ξ_j_*(*t*)〉 = *σ*^2^*δ_ij_δ*(*t* – *t*), and *i,j* = {1, 2, *c*}. Therefore, each neuron receives drive from one independent noise source, and one common noise source. The correlation in the drive between the two noise sources is *c*, and the input to neuron 2 is lagged by an amount *D*. Given this input, we can calculate the self-consistent dynamics for the synaptic weights, as before, assuming that the impact of each spike pair is weak compared to the dynamic range of the synapse. The equations are

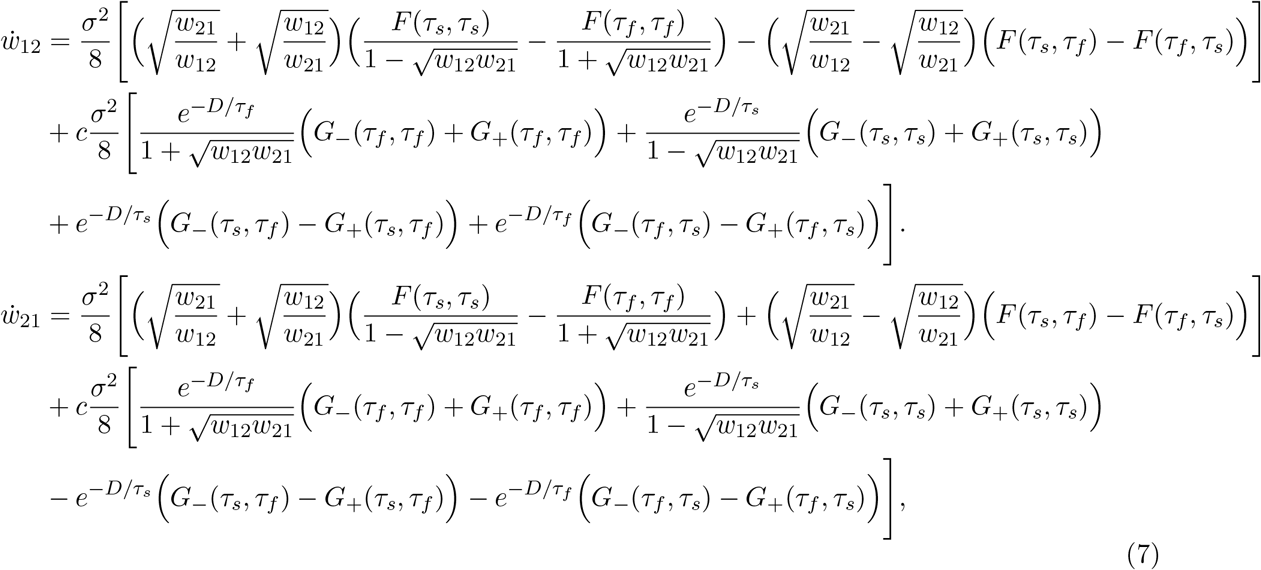

where

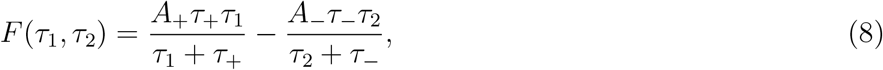

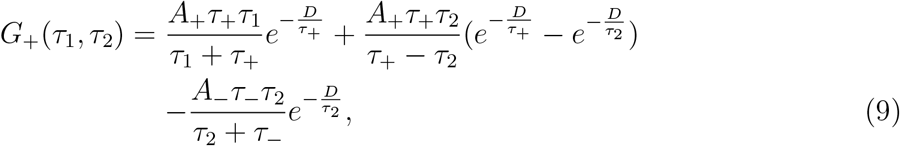

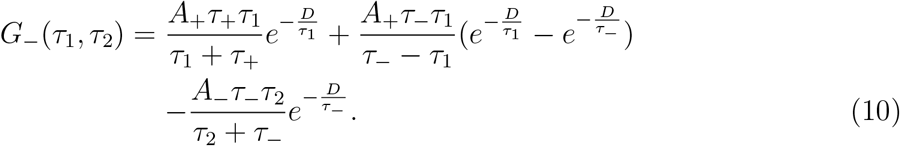

Note that lim_*D*→0_ *G*_+_ = lim_*D*→0_ *G*_ − = *F*. It may appear from Eqs.7 that the dynamics is singular for *w*_12_ → 0 or *w*_21_ → 0, but these limits are, in fact, well defined, see *Appendix C* for details.

As before, there are no fixed point solutions of Eqs.7. However, the equations together with the condition that 0 ≤ *w_ij_* ≤ *w_max_*, can be used to determine which synaptic states are stable, as in the previous section. Fig.6 shows sample phase planes as a function of the correlation and delay in the common noisy drive, and for three values of *w_max_*. In all cases, the fully potentiated state (PP) and the asymmetric state (DP) are favored for positive correlations. The state (DP) corresponds to a potentiated connection from 1 to 2, which occurs when the delay is sufficiently long (2 follows 1 here). For negative correlations the phase diagram is more complex, allowing for up to eight distinct regions. In general, the fully depressed state is stable when correlations are negative and the delay is not too large. On the other hand, the asymmetric state (PD) is stable everywhere for negative *c*.

**FIG. 6.**
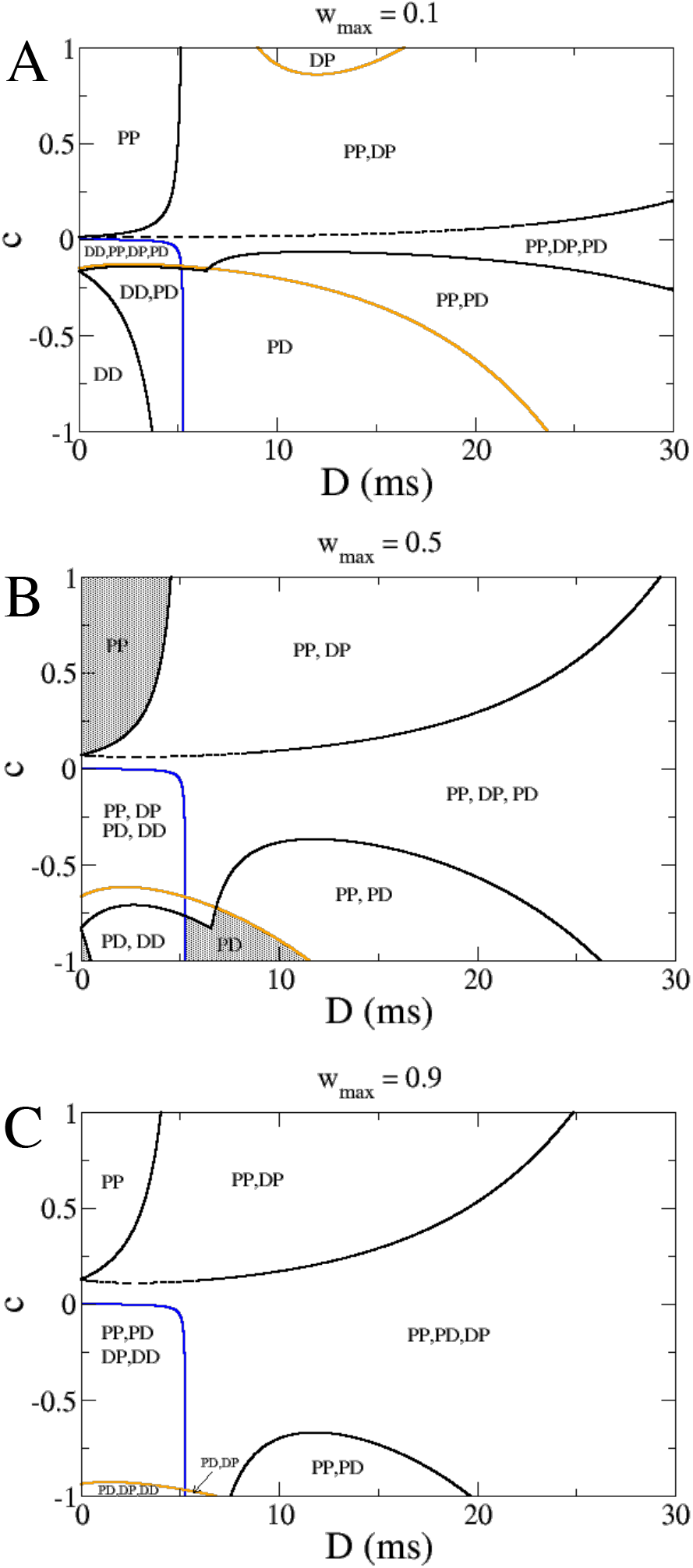
Phase planes of synaptic states for two noise-driven neurons as a function of the correlation and delay in the shared noisy drive. A. Phase plane for the case *w_max_* = 0.1. B. Phase plane for the case *w_max_* = 0.5. C. Phase plane for the case *w_max_* = 0.9. Other parameters: *τ* = 10ms, *σ* = 5, *I*_0_ = 10, *A*_+_ = 0.001, *τ*_+_ = 20ms, *τ*_−_ = 60ms, *A*_−_ = *A*_+_*τ*_+_/*τ*_−_. The shaded regions in B. indicate a single stable state.

Fig.7 shows a detail of the phase diagram in Fig.6B with symbols indicating parameter values use to confirm the analytical results through several illustrative numerical simulations.

**FIG. 7.**
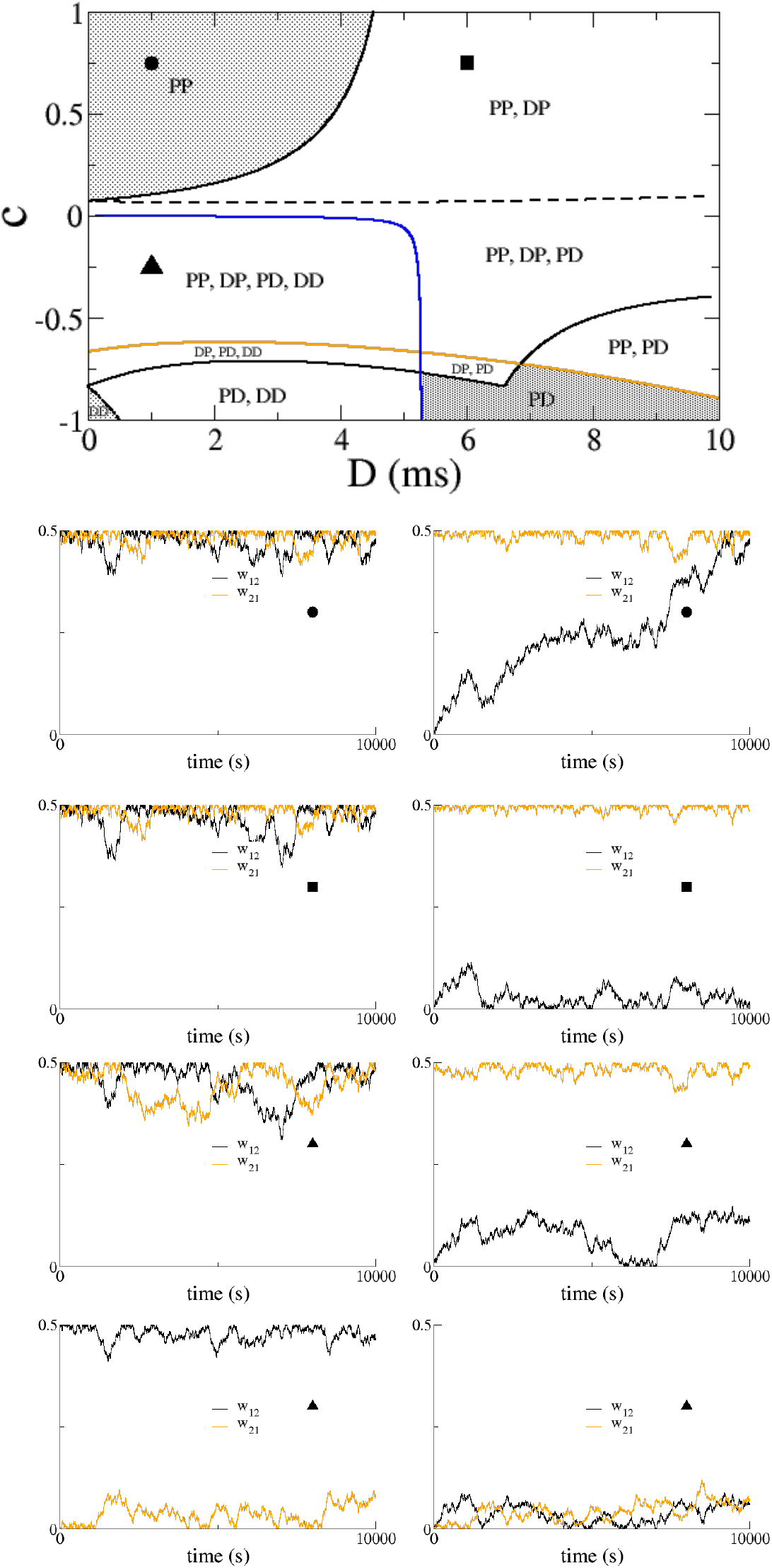
A blow-up of the phase plane from Fig.6B, with illustrative numerical simluations. The symbols indicate parameter values used for the numerical simulations.

## V. DISCUSSION

We have derived a set of equations for the joint evolution of neuronal activity and of the synaptic weights between pairs of neurons assuming a separation of time-scales between the two processes, Eqs.4. The resulting equations can be solved analytically in the case of linear firing rate neurons, and reduce to a set of coupled ODEs for the synaptic weights alone. For periodic and noisy forcing the resulting equations are Eqs.5 and Eqs.7 respectively. The plasticity rule we have chosen is formally a spike-timing dependent rule (STDP), although the fact that we generate spikes as a Poisson process insures that the actual spike timing plays no role here. Rather it is only variations in the underlying rates for the Poisson processes which can lead to plasticity in our model. This appears to be the dominant factor in shaping plasticity given realistic spike trains, even when spiking correlations are taken into account [6]. We additionally assume that the plasticity rule is balanced, namely that the integral over the STDP window is identically zero. If this is not the case, then the synaptic weights between neurons with non-zero rates would always grow or decay, depending on the sign of the integral. This implies that in network simulations the final synaptic weight matrix will always saturate or decay to zero, ruling out the emergence of any non-trivial structure. Additional mechanisms, such as homeostasis, are needed in this case in order to avoid saturation [18]. On the other hand, with the balance assumption only time-variations in the firing rates can drive plasticity. Specifically, the change in a given synaptic weight depends on the covariance of the pre- and post-synaptic firing weights, multiplied by the STDP window [14].

We also note that our equations do not take into account the variability arising due to the stochasticity of firing, but rather only the mean Poisson rates. In numerical simulations of the full, stochastic system, one observes fluctuations which may momentarily drive the synaptic weight away from its stable meanfield value (this is clearly seen in Fig.7) or even cause transitions between stable states. These effects are not captured by Eqs.5 or Eqs.7.

### Oscillatory drive

In the case of two periodically forced neurons, the resulting connectivity motif depends solely on the phase difference, while the time it takes for the connectivity to reach its steady state depends strongly on the forcing frequency. When potentiation dominates at short latencies then small phase differences lead to a fully potentiated motif. Larger phase differences generate a unidirectional motif in which the connection from leader to follower potentiates whereas the other depresses. Finally in the vicinity of anti-phase forcing both synapses depress and the neurons decouple. Analysis of Eqs.5 furthermore reveals several regions of bistability between these different motifs. Simulations agree well with the analysis, see Fig.3. Previous work on STDP in a network model of hippocampus found similar effects of the phase of oscillation on connectivity motifs through simulation [3].

The evolution equations for the synaptic weights of a network of arbitrary size can be derived using the separation of time scales technique. However, for several cases of interest the theory for pairs of neurons can be used to gain insight into the resutling connectivity of large networks. This includes the case of several clusters forced at different phases, as well as the case of a uniform distribution of phases. When phases are widely distributed the general finding is the emergence of hierarchical structure. Specifically, there is clustering locally between neurons with similar phases of the forcing, but between neuron pairs with disparate phases, unidirectional connections form according to the leader-follower phase-relationship. It is interesting to note that precisely this type of hierarchical clustering in cortical microcircuits has been inferred from data collected through multiple patch-clamp experiments in slices [16].

### Noisy drive

In the case of two neurons forced by white-noise inputs with a given correlation and time-delay, the resulting phase diagram is very rich, see Fig.6. Although positive correlations tend to lead to potentiation and negative correlations to depression, the combined effect of correlation and delay is complex, and multistability is the rule. This is borne out in numerical simulations for pairs of neurons, see Fig.7. It is unclear how this will affect the emergent connectivity in large networks of neurons, and requires additional study.

The plasticity process we describe here consists of a build up over time of a large number of small changes. As such, it is slow, and would not be a relevant mechanism for rapid memory formation, such as episodic memory. Rather, such a process shapes the connectivity in recurrent circuits in accordance with regularities in the statistics of the inputs. For example, if we consider an area in the visual pathway, neurons with similar feature selectivity and overlapping receptive fields would exhibit positive correlations in their output in response to a time-varying stimulus. Both for the case of oscillatory as well as noisy dynamics our analysis would predict a potentiation of recurrent connections between these neurons. This would lead to an enhanced response to a similar stimulus over time. Neurons with similar feature selectivity but non-overlapping receptive fields would likely exhibit similar yet time-delayed (or out-of-phase) inputs in response to the motion of an object across the visual field, etc. Therefore, we expect that the statistics of some sensory stimuli can be mapped on to the parameters of the inputs in our model: frequency, phase, input-correlation, delay. Repeated exposure to the same sensory stimuli would lead to a slow reshaping of the recurrent circuit connectivity and hence the neuronal response. Such a process may be relevant for the phenomenon of perceptual learning [4].

## APPENDIX A: DERIVATION OF SELF-CONSISTENT EQUATIONS FOR SYNAPTIC WEIGHTS

Here we derive a self-consistent set of equations for the synaptic weights *w*_12_ and *w*_21_ by assuming a separation of time-scales between the rate dynamics and the synaptic plasticity. We first note that the evolution equations for the synaptic weights, Eqs.2 can be rewritten for the case of stationary rate dynamics by noting that 〈*r_i_*(*t*)*r_j_*(*t* – *T*)〉_*t*_ = (*r_i_*(*t*+*T*)*r_j_*(*t*))_*t*_, which allows for a change of variables in the second integral, leading to Eq.3. Strictly speaking this correspondence only holds when the dynamics has been averaged over the fast time. As we are only interested in the slow-time dynamics, we write Eq.3 as if the correspondence were exact, cognizant that it is a slight abuse of notation.

Now, given real-valued, time-varying inputs to the two neurons, the equations are

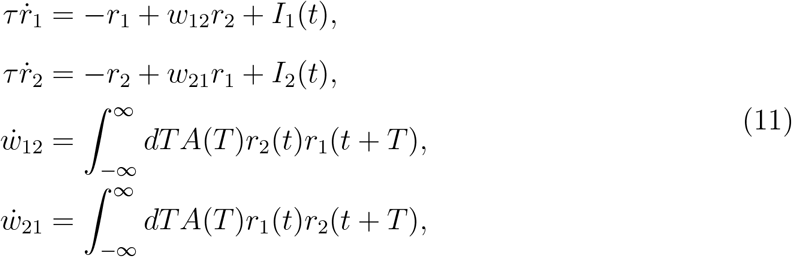

where *A*(*T*) is the plasticity rule. There is no general analytical solution to these equations given the quadratic nonlinearities. However, if we assume that each synaptic weight change is small, then the synaptic weights will evolve much more slowly than the rates and we can formally separate the time scales in a multi-scale analysis. To do this we introduce the small parameter *ϵ* ≪ 1 such that *A*(*T*) = *ϵÃ*(*T*). We also introduce the slow time *t_s_* = *ϵt* and allow for the rates and weights to evolve on both fast and slow time scales, i.e. they are functions of *t* and *t_s_*, and these two times are taken to be independent variables.

Then we can write

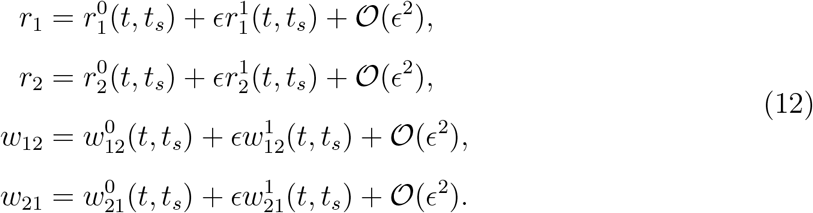

Pluggin these into Eqs.11 and collecting terms order-by-order gives, at order 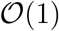

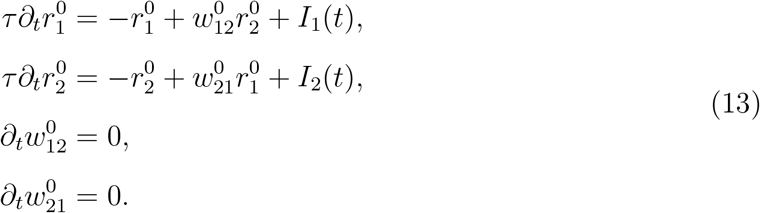

These last two equations show that the leading-order weights only depend on the slow time, namely 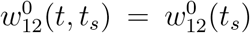 and 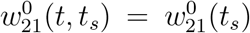. Therefore, they can be treated as constants in the rate equations, which evolve on the fast time-scale.

At order 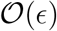 we have

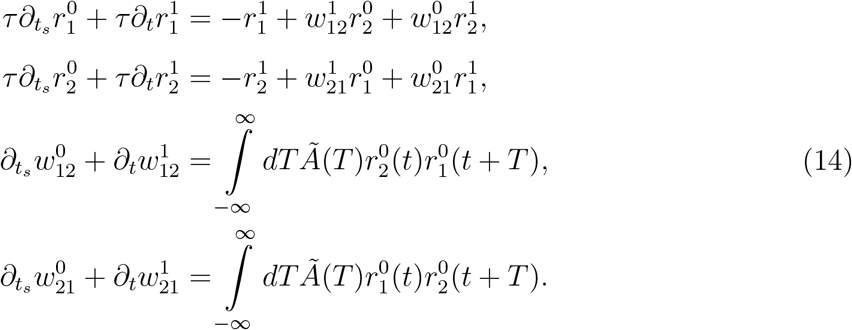

The first two equations give a correction to the leading order solution of the firing rates, which we will not use here. The weight equations at first glance do not seem solvable since we are expected to solve for both the leading order solution of the synaptic weights as well as the next order correction in the same set of equations. However, we know that the leading order terms are independent of the fast time *t*, which will allow us to solve for both. Specifically, the evolution of the leading-order weights will depend only on those terms from the integral which are independent of the fast time. This leads to Eqs.15. For simplicity in notation, in what follows we will drop the superscripts and tildes and write, for the leading-order solution, simply

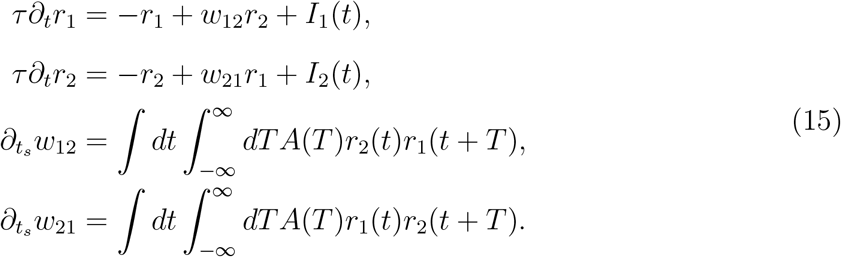

We will consider the specific cases of oscillatory and noisy drive below.

## APPENDIX B: OSCILLATORY DRIVE

Here we study the case where the neurons are driven sinusoidally with a frequency *ω* and with a phase difference of *ϕ*, i.e. *I*_1_ = *I*_0_ + *Ie*^*iωt*+*iϕ*_1_^ and *I*_2_ = *I*_0_ + *Ie*^*iωt*+*iϕ*_2_^. The (complex) rates can be written (*r*_1_, *r*_2_) = (*R*_10_(*t_s_*), *R*_20_(*t_s_*)) + (*R*_11_(*t_s_*), *R*_21_(*t_s_*))*e^iωt^*. We find that

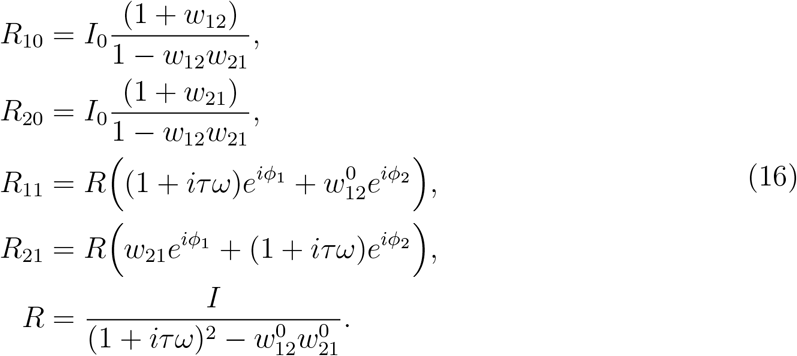

Because we only consider balanced plasticity rules here, the constant, baseline rates will not affect the synaptic rates. Nonetheless, when conducting numerical simulations it is important to take large enough constant drive *I*_0_ to ensure positive rates.

In order to calculate the equations for the synaptic weights we must use the real part of the complex rates. As an illustration we consider the equation for the weight *w*_12_, which is

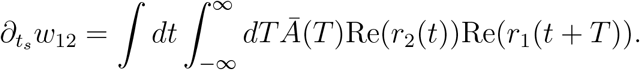

The quadratic term

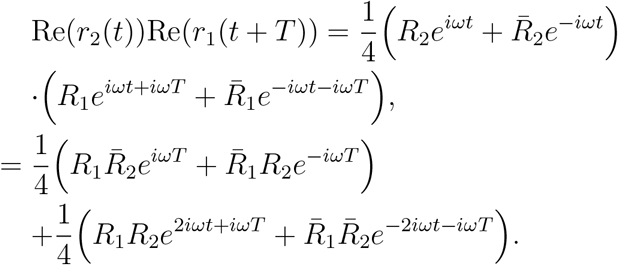

Note that the first two terms are independent of the fast time *t*, while the second two terms oscillate on the fast timescale with a frequency 2*ω*. Integrating over the fast timescale therefore eliminates the latter terms. Performing the second integral and then doing the analogous calculation for the other weight leads to Eqs.5 where *φ* = *φ*_2_ – *φ*_1_.

### Growth rate of synaptic weights

While the final state of the synaptic weights depends on the phase difference, the rate at which plasticity occurs is strongly influenced by the frequency of forcing. This can be most easily seen for the case of in-phase forcing *ϕ* = 0, for which we expect both weights to potentiate (or depress for an anti-hebbian rule). Assuming *w_ij_* = *w_ji_* leads to a right-hand side (growth rate) of Eqs.5 which is simply proportional to *Ã*_+_(*ω*) – *Ã*_−_(*ω*) which is zero for *ω* = 0 and as *ω* → ∞, while it has a maximum for 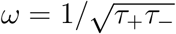.

#### Theory for networks

For the case of *n* coupled neurons, the rate equation for the ith neuron is

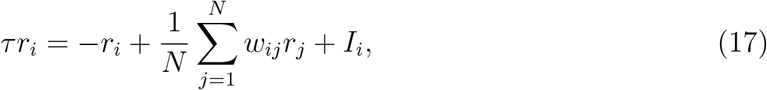

where *I_i_* = *I*_0_ + *Ie*^*iϕ_i_*^, while the evolution equation for the synaptic weight from neuron *j* to neuron *i* is still described by Eq.3. We can once again apply the separation of timescales formally by defining *A*(*T*) = *ϵÃ*(*T*) where *ϵ* ≪ 1 and defining the slow time *t_s_* = *ϵt*. The rates can be written in vector form as **r**(*t,t_s_*) = **R**_0_(*t_s_*) + **R**_1_(*t_s_, ω*)*e^iωt^* where

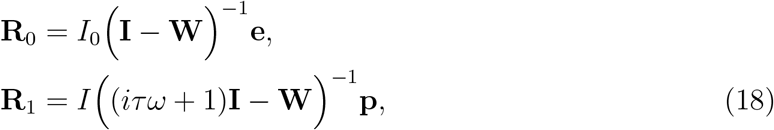

where **I**, **W** are the identity matrix and weight matrix respectively, **e** is a vector of ones, while the *j*th element of the vector **p** is *e^iϕj^*.

### Applicability of pairwise theory to network simulations

If we consider the rate equations for a pair of neurons *j* and *k* in the network Eqs.17 we find, applying the separation of time-scales approach detailed in Appendix A, that the oscillatory components obey

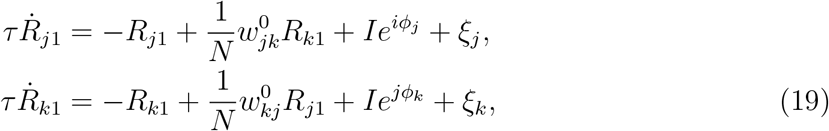

where 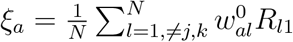. From this we find the complex amplitudes

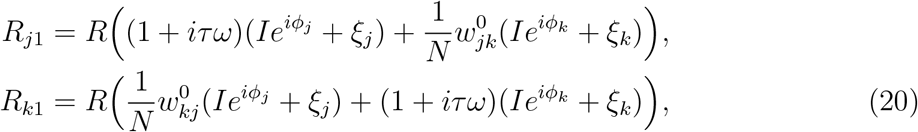

where 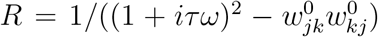. Note that these equations are identical to those for the complex amplitudes for the pairwise case (with renormalized weights), Eqs.16, with the exception of the meanfield terms *ξ_j_* and *ξ_k_*. The slow dynamics of the synaptic weight *w_jk_* is then given by

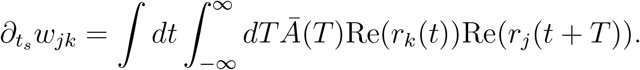

Note that in principle the rates *r_k_* and *r_j_* still depend on the meanfield terms and hence this equation is not self-consistent as in the pairwise case. The influence of these meanfield terms depends strongly on the distribution of phases of the complex amplitudes. In one of the two limiting cases, if all of the phases are aligned then the moduli of the terms all sum. This is equivalent to the summation of vectors all with the same angle. In the other limiting case, if the phases are uniformaly distributed, then the resultant modulus will be close to zero because we are summing many vectors all with distinct phases (as long as the moduli and phases are only weakly correlated or uncorrelated). Hence in this limit the inlfuence of the meanfield vanishes and only the pairwise interactions matter. This latter case is the relevant one for Fig.5C and explains why the pairwise theory correctly predicts the network structure after learning.

### Network simulations

For the simulations shown in Fig.4, the following nonlinear rate equations were used

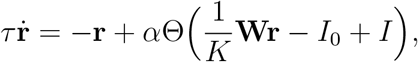

where Θ is the Heaviside function. A spike from a neuron *i* in a timestep Δ*t* occurs with probability *r_i_*Δ*t*. Given the spike trains from neurons *i* and *j*, a weight *w_ij_* undergoes updates from all spike pairs according to the STDP rule, see [11] for the numerical scheme. For the simulations in Fig.5 linear rate equations are used.

## APPENDIX C: NOISY DRIVE

Here we consider an external drive of the form

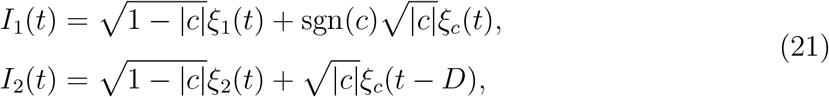

where *ξ_i_*(*t*) is a Gaussian white noise process, i.e. 〈*ξ_i_*(*t*)〉 = 0 and 〈*ξ_i_*(*t*)*ξ_j_*(*t*)〉 = *σ*^2^*δ_ij_δ*(*t* – *t*). Therefore, the noisy drive to the two neurons has correlation *c*. The correlated input is delayed to neuron 2 with respect to neuron 1 by a time *D*.

To solve the system of rate equations we rewrite it in vector form as

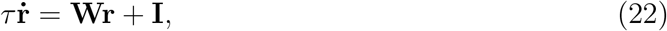

where **r** = (*r*_1_, *r*_2_), **I** = (*I*_1_, *I*_2_) and

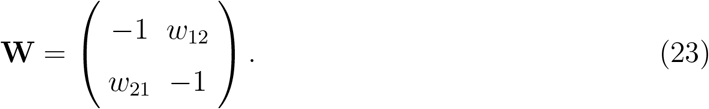

We diagonalize the connectivity matrix **W** = **QΛQ**^−1^ and obtain the system of independent equations

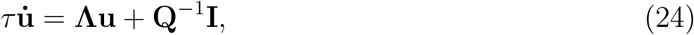

where **u** = **Q**^−1^**r**. The matrices resulting from the diagonalization are

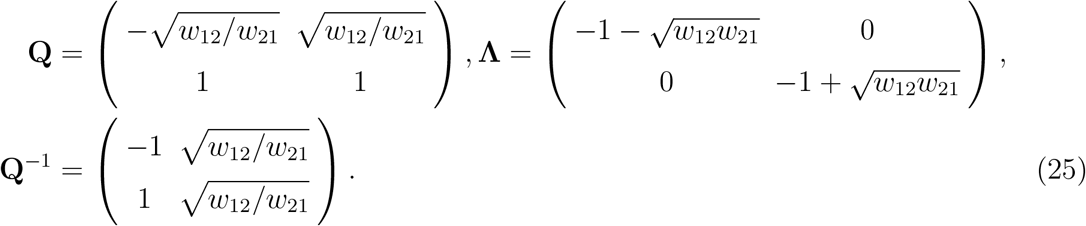

The equations for the transformed variables **u** are

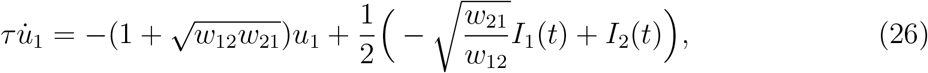

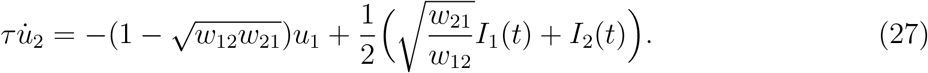

These equations can be solved formally as

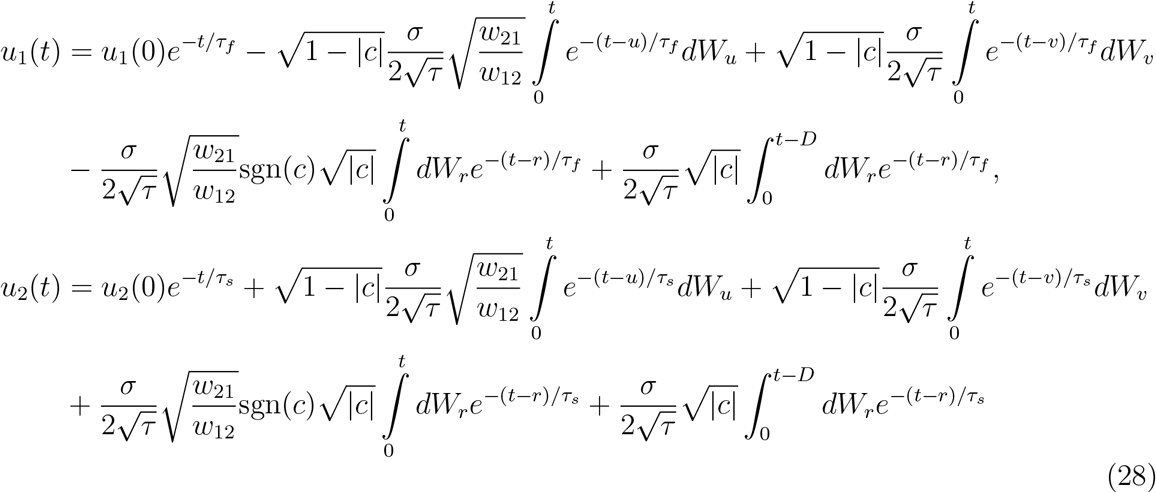

where *dW_u_, dW_v_* and *dW_r_* are the stochastic differentials corresponding to the Gaussian processes *ξ*_1_, *ξ*_2_ and *ξ_c_* respectively. Also, we have defined the fast and slow time constants

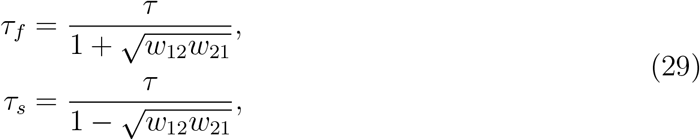

from which it is clear that there is an instability for *w*_12_*w*_21_ > 1. The original firing rates are linear combinations of these variables. Specifically,

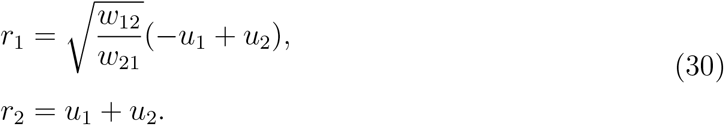

Finally, we have, and ignoring the dependence on the initial condition,

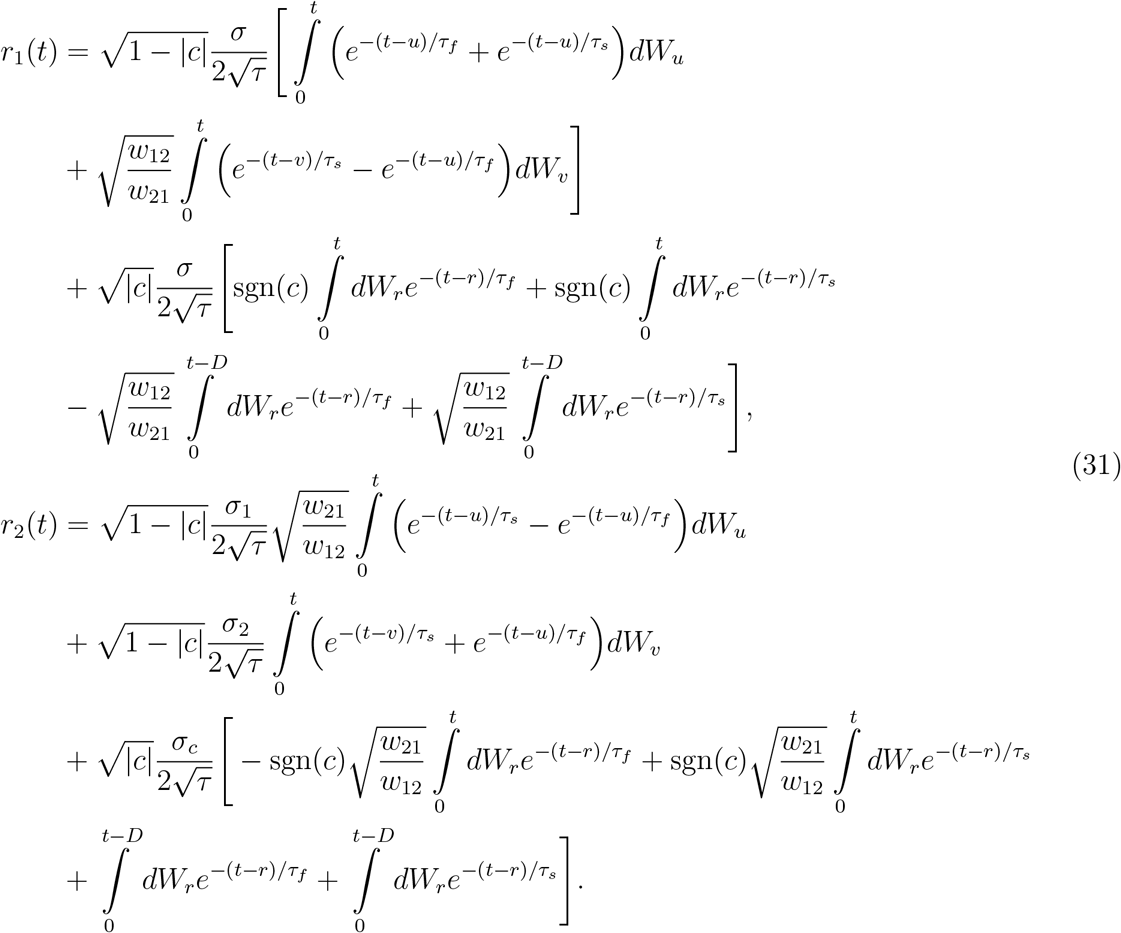

The slow dynamics of the synaptic weights, which is calculated self-consistently through the rates, is therefore also stochastic. In this case the integral over the fast time in Eqs.15 yields the expected value of the product of rates. Namely,

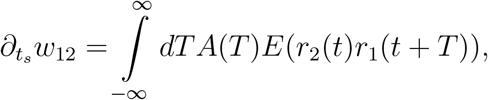

and similarly for *w*_21_. Evaluating this expectation requires products of stochastic integrals. For independent processes this expectation is always zero, while for integrals of the same process the product can be expressed as a standard integral through the so-called Ito isometry.

For example,

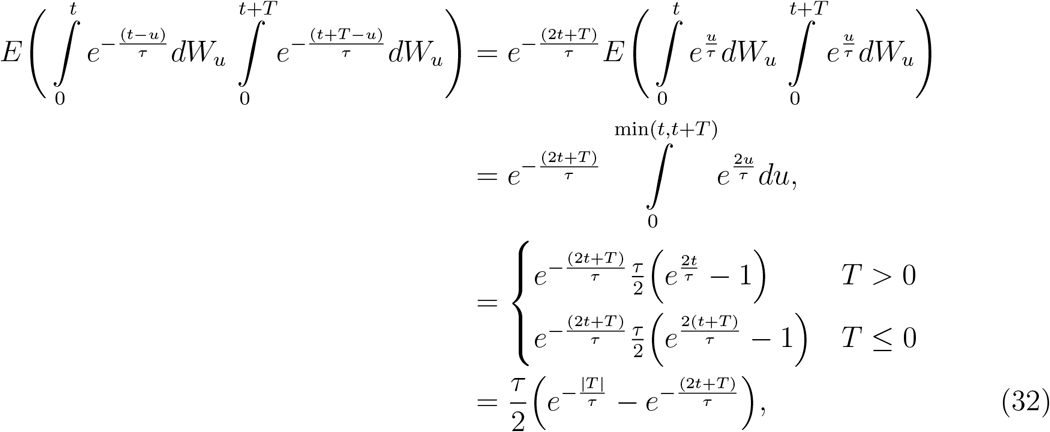

which is independent of *t* at long times. Performing these integrals yields the evolution equations, Eqs.7.

### Evolution equations for small weights

If *w*_12_ ≪ 1 and *w*_21_ ≪ 1 then

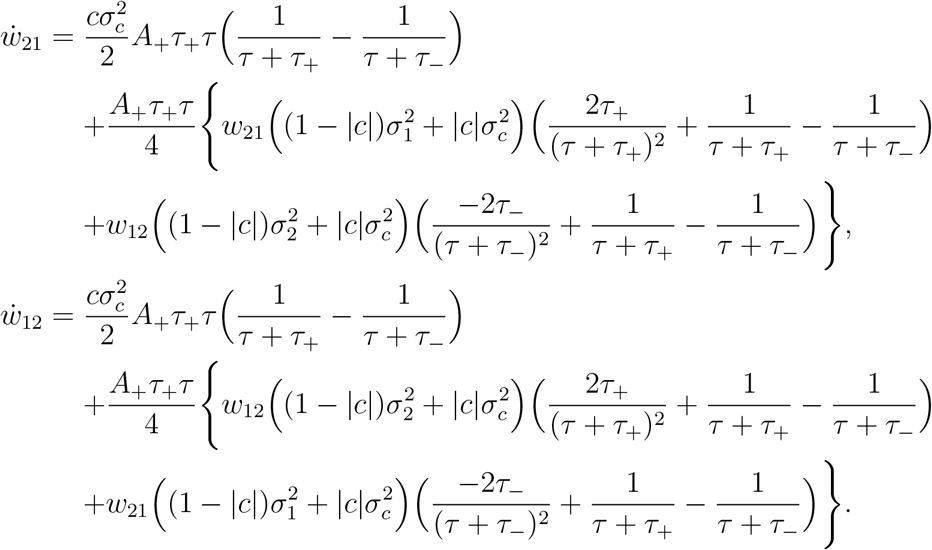

If *w*_12_ ≪ 1 and *w*_21_ can be order one, then

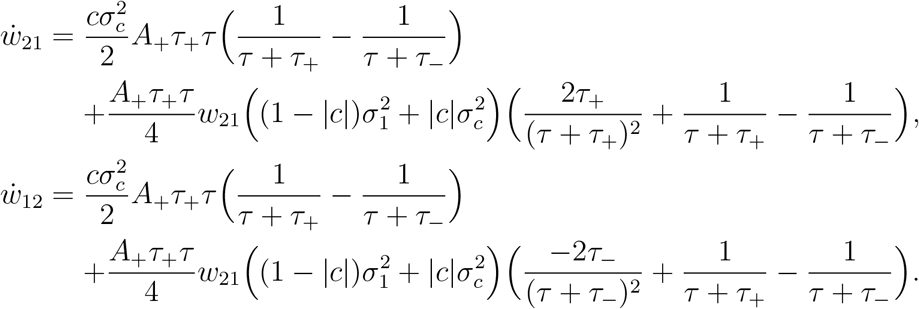

If *w*_21_ ≪ 1 and *w*_12_ can be order one, then

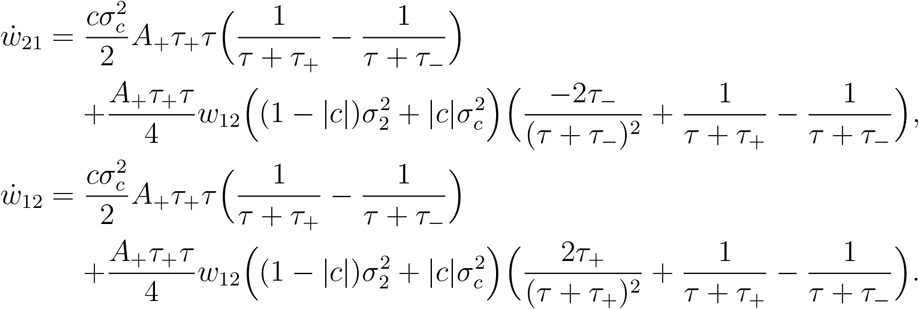

## ACKNOWLEDGMENTS

AR acknowledges “Retos” project RTI2018-097570-B-100 from the Ministry of Science and Innovation of the Spanish Government, Flag-Era project from the EU for the Human Brain Project HIPPOPLAST (Era-ICT code PCI2018-093095), “Red de Invetigación” RED2018-102323-T from the Ministry of Science and Innovation of the Spanish Government. This work is supported by the Spanish State Research Agency, through the Severo Ochoa and Maria de Maeztu program for Centers and Units of Excellence in R&D (CEX2020-001084-M). We thank CERCA Program/Generalitat de Catalunya for institutional support. We acknowledge very helpful discussions with Marina Veguó, Toni Guillamon and Ernest Montrbrióo.

